# Caveolae and scaffold detection from single molecule localization microscopy data using deep learning

**DOI:** 10.1101/526327

**Authors:** Ismail M. Khater, Stephane T. Aroca-Ouellette, Fanrui Meng, Ivan Robert Nabi, Ghassan Hamarneh

**Affiliations:** Medical Image Analysis Lab, School of Computing Science, Simon Fraser University, Burnaby, BC V5A 1S6, Canada; Department of Cellular and Physiological Sciences, LSI Imaging, Life Sciences Institute, University of British Columbia, Vancouver, BC V6T 1Z3, Canada

## Abstract

Caveolae are plasma membrane invaginations whose formation requires caveolin-1 (Cav1), the adaptor protein polymerase I, and the transcript release factor (PTRF or CAVIN1). Caveolae have an important role in cell functioning, signaling, and disease. In the absence of CAVIN1/PTRF, Cav1 forms non-caveolar membrane domains called scaffolds. In this work, we train machine learning models to automatically distinguish between caveolae and scaffolds from single molecule localization microscopy (SMLM) data. We apply machine learning algorithms to discriminate biological structures from SMLM data. Our work is the first that is leveraging machine learning approaches (including deep learning models) to automatically identifying biological structures from SMLM data. In particular, we develop and compare three binary classification methods to identify whether or not a given 3D cluster of Cav1 proteins is a caveolae. The first uses a random forest classifier applied to 28 hand-crafted/designed features, the second uses a convolutional neural net (CNN) applied to a projection of the point clouds onto three planes, and the third uses a PointNet model, a recent development that can directly take point clouds as its input. We validate our methods on a dataset of super-resolution microscopy images of PC3 prostate cancer cells labeled for Cav1. Specifically, we have images from two cell populations: 10 PC3 and 10 CAVIN1/PTRF-transfected PC3 cells (PC3-PTRF cells) that form caveolae. We obtained a balanced set of 1714 different cellular structures. Our results show that both the random forest on hand-designed features and the deep learning approach achieve high accuracy in distinguishing the intrinsic features of the caveolae and non-caveolae biological structures. More specifically, both random forest and deep CNN classifiers achieve classification accuracy reaching 94% on our test set, while the PointNet model only reached 83% accuracy. We also discuss the pros and cons of the different approaches.

## Introduction

Caveolae are tiny structures of 50–100 nm plasma membrane invaginations [1], membrane-attached vesicles, that have roles in membrane trafficking and signaling [2]. Caveolin-1 (Cav1) is the coat protein for caveolae, however formation of invaginated caveolae also requires the coat protein CAVIN1/PTRF. In the absence of CAVIN1/PTRF, Cav1 forms flat scaffold domains that have distinct functions from caveolae [3]. Secretion and overexpression of Cav1 in prostate cancer promotes tumor growth and has significant role in cancer metastasis [2]. Cav1 domains are below the diffraction limit of the light microscopy (i.e. 250 nm) which makes it hard to study them using conventional microscopic imaging modalities. Recent advancements in microscopy technology have enabled light microscopes to break Abbe’s diffraction limit. These techniques, known as super-resolution microscopy, can reach resolutions of < 20 nm in localizing the target protein [4]. Single molecule localization microscopy (SMLM) is a subset of techniques that work by manipulating the environment such that in each captured instance, a frame, only a few molecules are stochastically activated to emit light. Highly precise localizations can then be obtained from isolated point spread functions (PSFs) of isolated fluorophores (blinks). A 2D super-resolution image can be obtained by stacking up thousands of the collected frames. To achieve a 3D SMLM image, a cylindrical lens is inserted so that the microscope captures a deformed Gaussian PSF for each molecule. The XY coordinates of the molecule are measured as the center of the PSF, while Z coordinate can be measured from the deformation of the PSF [4, 5]. Consequently, the nanoscale 3D biological clusters with dimensions below the diffraction limit of optical light (i.e. 200–250 nm) can be studied and visualized using the final 3D point cloud collected from the SMLM frames.

Stone et al. [6] have applied super-resolution imaging to study the mammalian plasma membrane structure and organization. Sherman [7] reviewed how SMLM helped in studying the organization of signalling complexes in intact T cells. He concluded that the cell membrane employs dynamic and hierarchical patterns of interacting molecular species that have a critical role in cell decision making. Baddeley [8] studied the super-resolved SMLM techniques that are capable of examining biological structures in the cell membrane. He concluded that SMLM imaging methods are attractive techniques for investigating the proteins and receptors clustering. Khater et al. [9] and Baddeley [8] focused on the need for new computational tools for quantitatively analyzing the SMLM data. Khater et al. [10] studied the cellular structures in the membrane of the prostate cancer cells using super-resolution microscopy of single molecules. They proposed graphlet and modularity based machine learning method to identify Cav1 domains and their biosignatures from super-resolution SMLM images [10, 11].

Deep learning is a type of machine learning technique that has attracted great attention in the past several years [12], as it relieves the algorithm developer from having to design features for a variety of prediction problems and is capable of achieving state of the art results in many application areas including medical imaging [13]. For the SMLM imaging modality, deep learning has been applied to PSF localization, i.e. estimating the X,Y,Z coordinates of the individual molecules in the fluorescent state from the raw event data collected by the microscope [14–16]. However, to the best of our knowledge, deep learning has yet to be applied to the subsequent (post-localization) analysis and quantification of the localization data, e.g. identifying the various biological structures.

SMLM analysis of Cav1 has previously been reported in zebrafish [17, 18]. Super-resolution microscopy enabled them to study the colocalization of Cav1 and CRFB1 clusters and their role in antiviral signalling [17]. SMLM has also been applied to study caveolae deformation in response to hypotonic shock [19]. In this work, we focus on the analysis of SMLM images of PC3 cancer cell labeled with antibodies to the membrane protein Cav1. Cav1 can be localized to invaginated caveolae or non-caveolar scaffolds [3]. The presence of the CAVIN1/PTRF protein, a Cav1 adaptor protein, is required for the creation of a caveola [1]. Caveolae have functional roles in the cell as mechanoprotective membrane buffers, mechanosensors, signaling hubs and endocytic transporters [20]. The role of scaffolds is less well-characterized, in large part due to difficulties distinguishing these two Cav1-positive membrane domains, but they have been specifically associated with regulation of receptor signaling and prostate cancer progression [21, 22]. The primary objective of our research is to identify whether a given Cav1-positive membrane structure is or is not a caveolae.

SMLM data is difficult for humans to visually inspect and manually analyze as the data is noisy and contains hundreds of thousands or millions of points representing complex cellular structures. As SMLM technology is a recent development, the majority of the published methods on SMLM are related to the image acquisition, with less published work about quantitative analyses from SMLM data. Among the SMLM quantification methods, many primarily investigate how to accurately segment 2D SMLM point clouds into clusters representing individual cellular structures. These cluster analysis methods currently rely on the extraction and analysis of a few primitive features (radius, density, number of points, etc.) to describe the 2D clusters as in Owen et al. [23, 24], where they applied Ripley’s functions to analyze the 2D clusters of super-resolution data. Beyond segmentation, some methods use the features to identify, group, and query of the different types of clusters. Lillemeier et al. [25] used the number of points per cluster and the cluster’s radius to compare between the clusters of two SMLM imaging techniques for two types of cells. Rossy et al. [26] extracted cluster features that capture the circularity, number of points, radius, and density of every cluster and then found simple statistics for each feature alone to compare more than two types of clusters. Pageon et al. [27] used the cluster density and diameter statistics to compare between two types of clusters. Caetano et al. [28] proposed an analytical tool that to extract cluster density, diameter, and size and then statistically compare different types of clusters based on these features. In the work of Rubin-Delanchy et al. [29], a simple statistic of each individual cluster feature was used to compare the clusters of two different types of cells. The primary features were the number of points, radius, and density, which were used to compare between two types of clusters. Levet et al. [30] proposed a software called SR-Tesseler that can be used to segment the 2D clusters and extract elementary features for them, but without training a system to identify them automatically. The software extracts four simple features for every 2D cluster. Their software is capable of extracting the area, number of points, circularity, and diameter of the individual clusters.

The aforementioned methods used a small number of features (cluster properties/descriptors) to quantify and analyze 2D (not 3D) SMLM clusters (blobs). The feature extraction methods used on 2D SMLM data are not sufficient to effectively identify and analyze these 3D clusters. Fortunately, the explosive growth in the field of machine learning over the last decade has yielded a number of algorithms that are able to analyze large data such as 3D SMLM data. In addition to being able to learn more and perhaps currently unknown features on its own, the machine learning approaches will also be capable to combine and weigh its learned features to automatically classify molecular structures. To our knowledge, we are the first to use machine learning to help in the identification and analysis of the SMLM data clusters.

In order to better understand the nature of the caveolae and its role in human biology, in this work, we have employed and compared a number of machine learning algorithms for identifying the caveolar structures from 3D SMLM data of PC3 cells.

## Materials and methods

### Methods overview

The primary objective of this research is to be able to accurately predict the class labels of segmented cellular structures originating from SMLM images of the same type of cells. We call these segmented structures blobs. We have approached this problem as a binary classification problem: caveolae (positive) or not caveolae (negative). Our approach to this problem involves three steps (described in detail later in the paper):

i. **Data pre-processing:** Denoises and segments blobs from SMLM data;
ii. **Data representation:** Describes the blob representations used (i.e., the representation of the input to the next step) we denote the transformation of the representation as *x* →*g*(*x*) = *x*′ where *x* is an input blob as a point cloud, *x*′ is a new representation of the same data; and *g* is the transformation function that may include transforming the point cloud into volumes, extract the 2D projections, etc.
iii. **Machine learning models:** Describes models used on each input representation and how they are trained to predict the class of a blob. We denote this prediction operation as *x*′*f* → (*x*′) = *f* (*g*(*x*)) = *ŷ* where *ŷ* is the predicted class (i.e. caveoalae or not). The function *f* is learned from a training set of *M* blobs with known class labels *{*(*x*_*i*_, *y*_*i*_), *i* = 1, 2, *…*, *M}*

### Image acquisition

PC3 prostate cancer cells (American Type Culture Collection (ATCC) and PC3 cells stable transfected with CAVIN1/PTRF-green fluorescent protein (GFP) (PC3-PTRF)(obtained from Michelle Hill, The University of Queensland Diamantina Institute, Brisbane, Australia) were cultured as previously described [1, 31] and plated on coverslips (NO. 1.5H, Carl Zeiss AG; coated with fibronectin) for 24 h before fixation with 3% paraformaldehyde (PFA) for 15 min at room temperature. Coverslips were rinsed with PBS/CM (phosphate buffered saline complemented with 1 mM MgCl2 and mM CaCl2), permeabilized with 0.2% Triton X-100 in PBS/CM, blocked with PBS/CM containing 10% goat serum (Sigma-Aldrich Inc.) and 1% bovine serum albumin (BSA, Sigma-Aldrich Inc.) and then incubated with the rabbit anti-caveolin-1 primary antibody (BD Transduction Labs Inc.) for 12 h at 4°C and with Alexa Fluor 647-conjugated goat anti-rabbit secondary antibody (Thermo-Fisher Scientific Inc.) for 1 h at room temperature. The primary and secondary antibodies were diluted in SSC (saline sodium citrate) buffer containing 1% BSA, 2% goat serum and 0.05% Triton X-100. Cells were washed extensively after each antibody incubation with SSC buffer containing 0.05% Triton X-100, post-fixed using 3% PFA for 15 min and washed with PBS/CM. Before imaging, cells were immersed in imaging buffer (freshly prepared 10% glucose (Sigma-Aldrich Inc.), 0.5 mg/ml glucose oxidase (Sigma-Aldrich Inc.), 40 *µ*g/mL catalase (Sigma-Aldrich Inc.), 50 mM Tris, 10 mM NaCl and 50 mM *β*-mercaptoethylamine (MEA; Sigma-Aldrich Inc.) in double-distilled water [4, 32] and sealed on a glass depression slide for imaging.

Ground state depletion microscopy (GSD) super-resolution imaging was performed on a Leica SR GSD 3D system using a 160x objective lens (HC PL APO 160x/1.43, oil immersion), a 642 nm laser line and an EMCCD camera (iXon Ultra, Andor). Preview images were taken with 5% laser power in both the GFP and Alexa Fluor 647 channels for each cell, in TIRF (total internal reflection fluorescence) mode. Full laser power was then applied to pump the fluorophores to the dark state; at a frame correlation value of 25% the imaging program auto-switched to acquisition with 50% laser power, at 6.43 ms/frame speed. The TIRF mode was also applied to the acquisition step of the GSD super-resolution imaging to eliminate background signals. The eventlist (i.e. SMLM data, also known as a point cloud) was generated using the Leica SR GSD 3D operation software with a XY pixel size of 20 nm, Z pixel size of 25 nm and Z acquisition range +/-400 nm. The CAVIN1/PTRF masks for the PC3-PTRF cells were generated by converting the GFP-channel of the preview images to binary images in ImageJ.

## Data

### Experimental data

The data used in this research comes from an experiment using PC3 prostate cancer cells [33]. The experiment is first run on 10 SMLM images from CAVIN1/PTRF absent PC3 cells, which from now on will simply be referred to as PC3 cells. It is then rerun on PC3 cells transfected with CAVIN1/PTRF-GFP, called PC3-PTRF cells (Fig 1). Due to imaging artifacts and high background signals, cell 6 of the PC3 cells and cell 7 of the PC3-PTRF cells were omitted from the data, leaving us with 9 PC3 and 9 PC3-PTRF cells. The experiment additionally captured lower resolution wide-field microscopy images of the GFP channel of PC3-PTRF cells to identify the location of CAVIN1/PTRF within each cell Fig 2. This mask provides us with a strong indication of where the caveolae are located and hence, we use it to label the blobs. Therefore, the blobs in PC3-PTRF data are labelled as PTRF-positive (PTRF+) and PTRF-negative (PTRF-). We used this mask and the known biology that caveolae contain more than 60 Cav1 molecules [9] to stratify the PTRF+ blobs into PTRF+≥60 and PTRF+< 60. Since caveolae cannot exist in PC3 cells, all blobs in PC3 cells were labeled as PTRF-negative (not caveolae or scaffold) as shown in the red color in Fig 1B.

**Fig 1.**
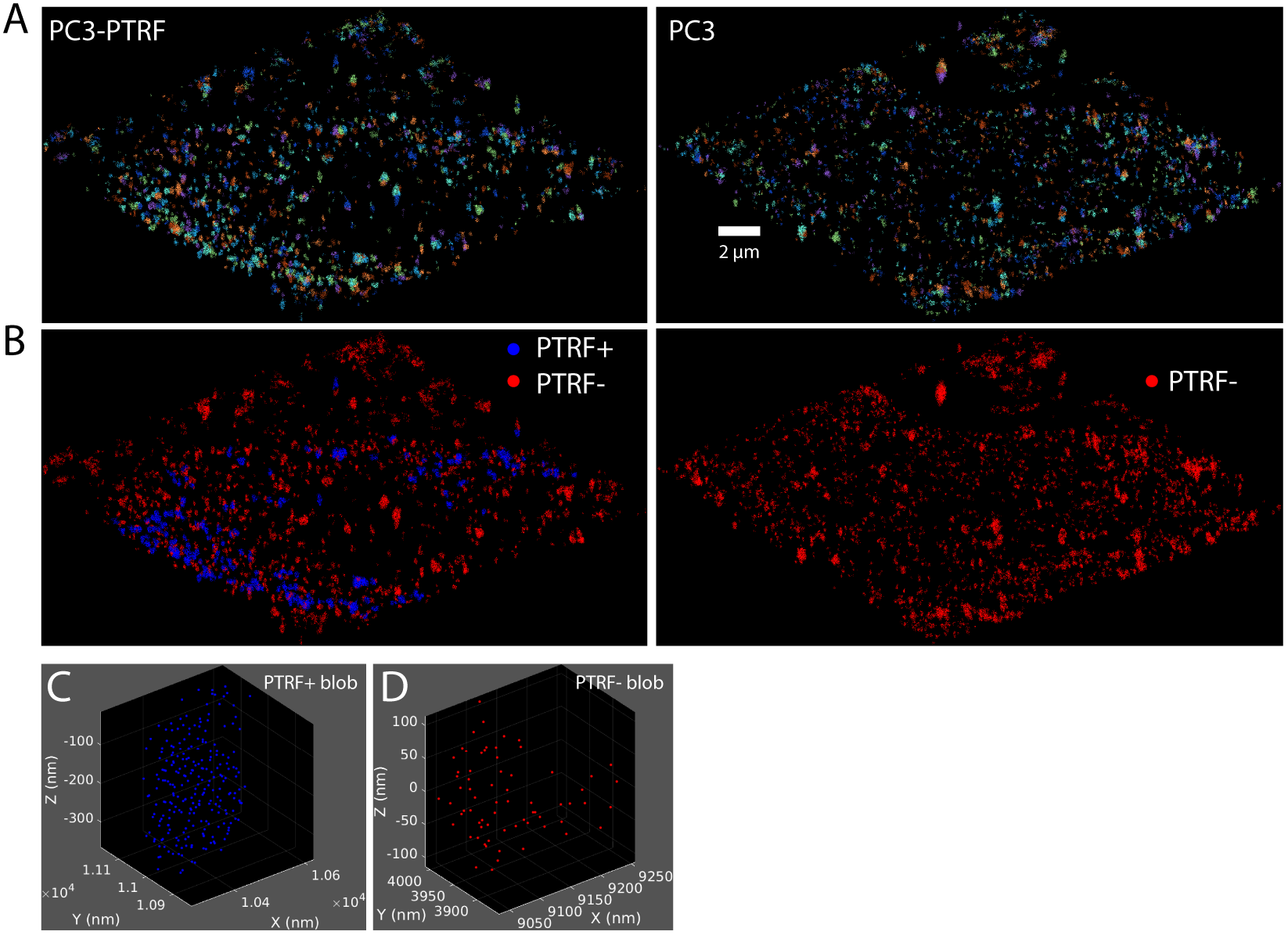
The process of obtaining Cav1 blobs (clusters) for the various learning tasks. Filtering, segmenting and labeling the blobs from the PC3 and CAVIN1/PTRF-transfected PC3 cells (PC3-PTRF cells). (A) 3D view of PC3 and PC3-PTRF cells of size 18×18×1 *µm*^3^. The view showing all the blobs (3D clusters) within a cell after applying the *3D SMLM Network Analysis* computational pipeline [9]. The pipeline contains modules to reconstruct the Cav1 molecules via the iterative merging of the localizations, filtering the noisy localizations, and segmenting the Cav1 blobs. The different colors show the segmented Cav1 blobs within the cell. (B) The blobs are color-labelled as PTRF-positive (PTRF+) and PTRF-negative (PTRF-). It shows that PC3 cell only has PTRF-blobs (non-caveolae blobs) that appear in red while the PC3-PTRF cell has both PTRF- and PTRF+ blobs that appear in red and blue respectively. (C) A sample PTRF+ blob taken from the PC3-PTRF cell showing the Cav1 molecules distributions. (D) A sample PTRF-blob taken from the PC3 cell showing the Cav1 molecules distributions.

**Fig 2.**
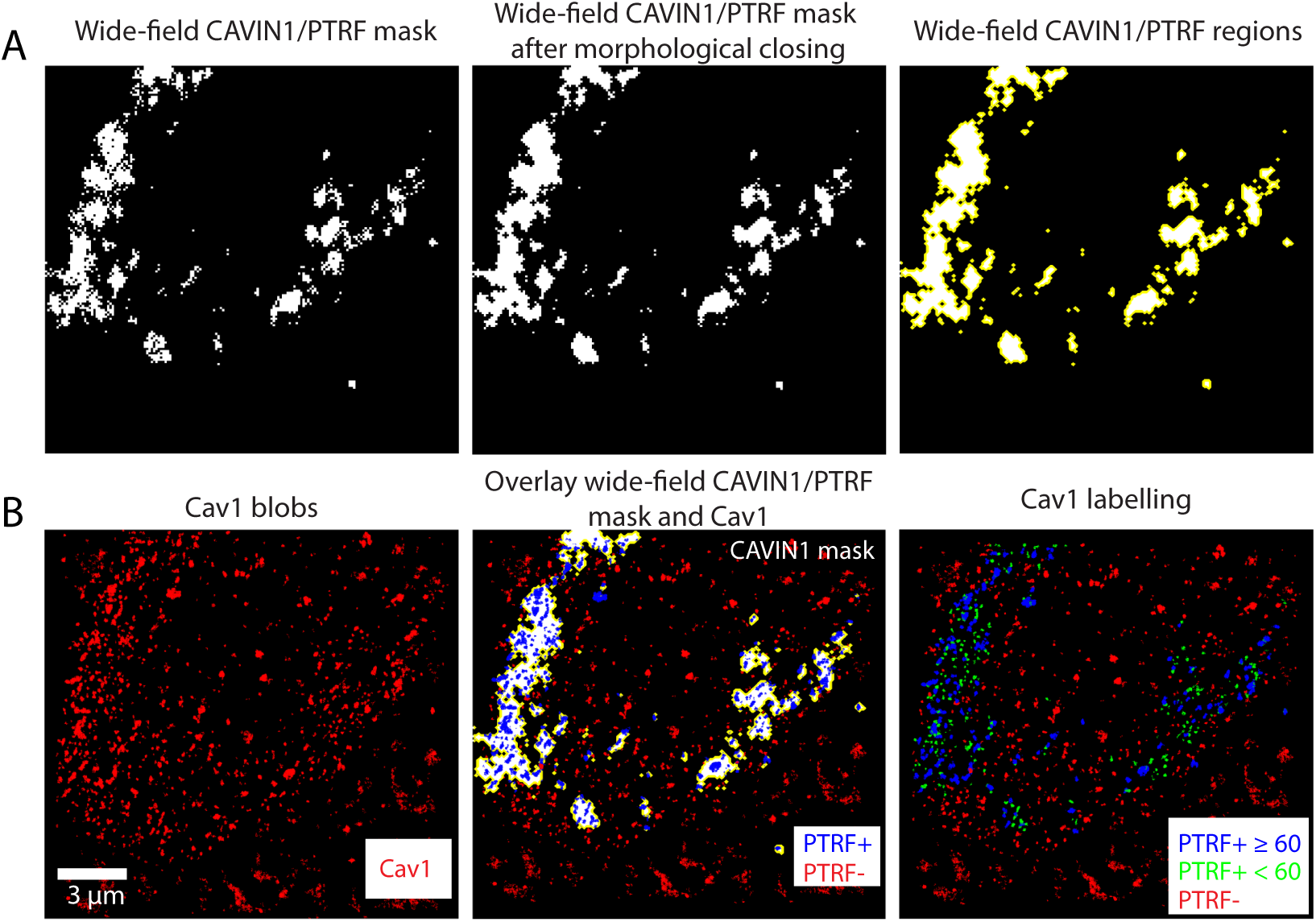
The process of obtaining the class labels for the Cav1 blobs using wide-field CAVIN1/PTRF mask. The class labels are necessary to train the machine learning models to identify the Cav1 blobs types automatically. (A) The first row shows the imaged wide-field TIRF CAVIN1/PTRF mask before and after morphological closing. The morphological closing operation is used to close the small holes in the consecutive regions of CAVIN1/PTRF mask. The CAVIN1/PTRF regions are delineated in yellow to highlight the locations of the CAVIN1/PTRF regions in the cell. (B) The second row shows the Cav1 blobs and the overlay of the Cav1 blobs with the wide-field CAVIN1/PTRF mask to label the blobs into PTRF+ and PTRF-. The caveolae structures have a minimum of 60 Cav1 molecule per blob [9] that can be used to stratify the PTRF+ blobs into PTRF+≥60 and PTRF+< 60. Our goal is to use machine learning approaches to automatically identify the PTRF+≥60 blobs (caveolar domains) from the rest of the non-caveolar domains (i.e. PTRF+< 60 and PTRF-) using different features and data representations of the blobs.

For our binary classification task, the 9 PC3 cells provide us 14491 negative blobs. The PC3-PTRF cells provide us 857 positive blobs (PTRF+≥60) and 10009 negative blobs (PTRF- and PTRF+< 60). To solve this data imbalance, we randomly downsample the negatives from 24500 blobs to 857 blobs to match the number of positives blobs. Fig 1B and and Fig 2 show the blobs from the two populations and their corresponding class labels before and after the number of molecules stratification respectively.

### Simulated data

Simulated data can help in validating the methods. We want to generate a simulated dataset of blobs with known class labels that mimic the real experimental dataset. In the real experiments, we are mainly studying two kinds of biological structures, i.e. caveolae and non-caveolar scaffolds. In our simulation, we are generating blobs that are similar to both classes. Specifically, we are generating a balanced dataset of 1000 blobs of isotropic point clouds and 1000 blobs of non-isotropic point clouds. The isotropic class of blobs mimicking the caveolae (positive class) and the non-isotropic class mimicking the non-caveolae (negative class). The non-isotropic class of blobs are more planar structures, while the isotropic class are more spherical structures. To simulate the real dataset, the number of points per generated blob is drawn randomly from 60–210 in the positive class and 10–160 in the negative class (Fig 3B). This insures that the blobs could have various number of points per blob in both classes. Also, the negative blobs might have a number of points that is equal or greater than the number of points in some of the positive blobs. Fig 3A shows two samples of the simulated dataset from both classes.

**Fig 3.**
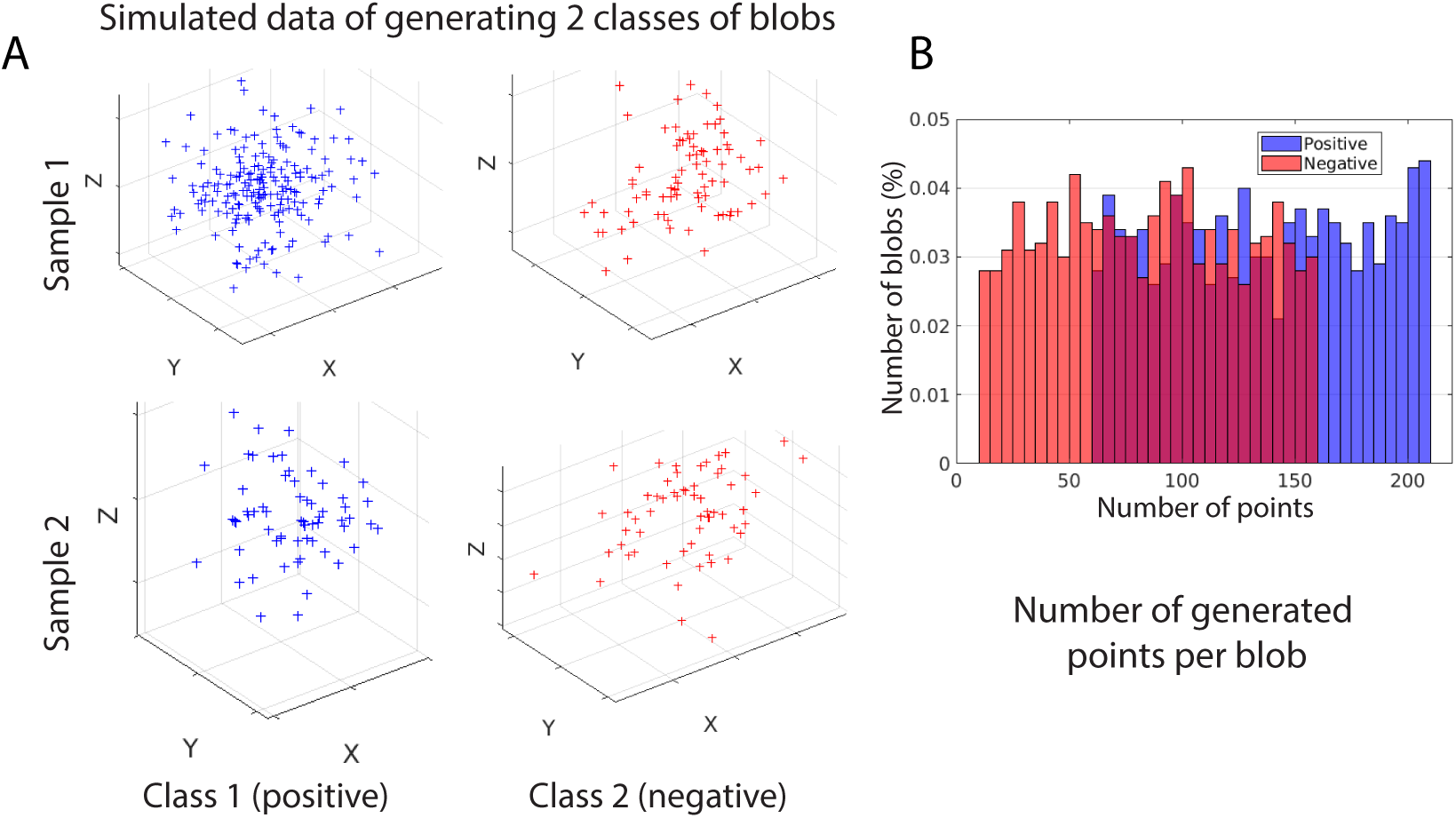
Simulated dataset that contains two types of simulated blobs that are generated with known classes. (A) The figure shows two samples from each class of the simulated data. The samples are taken from a dataset of 2000 blobs. The blue blobs are samples from the class 1 (positive) and the red blobs are samples from class 2 (negative). (B) The number of generated points per blob in both classes. It shows the percentage of generated blobs from both classes that have a variable number of points per blobs. There is a large overlap between many of blobs from both classes in terms of the number of generated points per blob.

In our simulation, we used the multivariate normal distribution to generate the samples of the two classes. Please see the following probability density function (pdf) of the 3-dimensional multivariate normal distribution that we adopted in our simulation experiments, equation 1.

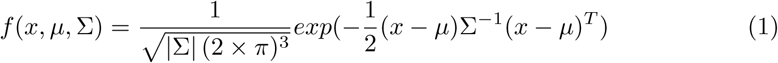

Where *x* and *µ* are 1×3 vectors and Σ
is a 3×3 symmetric, positive definite matrix. For the generated blobs from class 1 (isotropic), we used the covariance matrix 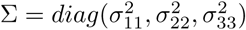, where the standard deviation *σ* = 10 nm. For the generated blobs from class 2 (non-isotropic), we used the covariance matrix 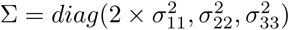, where the standard deviation *σ* = 10 nm. For both classes, we generated each blob to be centered at zero, i.e. *µ* = [0, 0, 0].

## Results and Discussion

### Data pre-processing

We adopted the computational pipeline of Khater et al. [9] to pre-process and post-process the SMLM data. The iterative merging algorithm is used for molecular reconstruction and correcting for multiple blinking of a single fluorophore artifact by iteratively merging all the localizations within 20 nm until converging to the predicted Cav1 localizations. The unclustered Cav1 molecules, as well as the background events, are removed via the filtering module by comparing the features of the Cav1 network with a random network. The clustered Cav1 node features are retained due to their distinct features as compared to the random network nodes features. Their pipeline then segments each cluster into individual cellular structures, i.e., blobs. Fig 1A shows two cells from both populations (PC3 and PC3-PTRF) after the denoising and segmentation of the blobs. Fig 1B shows the blob labelling of PC3-PTRF cell using the corresponding CAVIN1/PTRF mask that creates two types of blobs, PTRF-positive (PTRF+) that match with the mask and PTRF-negative (PTRF-) not-caveolae blobs outside the mask. The caveolae structures have a minimum of 60 Cav1 molecule per blob [9]. Therefore, in Fig 2 we show the PTRF+ blobs are stratified based on the number of molecules into PTRF+≥60 and PTRF+< 60. The PC3 cell expresses one type of blob in the absence of CAVIN1/PTRF protein. Hence, they mainly have one class label.

### Data representation

The application of the pre-processing pipeline results in a set of segmented blobs and their associated labels identifying them as caveolae (PTRF+≥60) or not-caveolae (PTRF- and PTRF+< 60) as seen in Figs 1C and D respectively. The blobs are left in the original point cloud format. While this representation has some benefits, it also has drawbacks and is not commonly used in deep learning. We, therefore, investigate a number of different input representations. Figs 4A–D shows the different representations a given blob can take for the different machine learning tasks.

**Fig 4.**
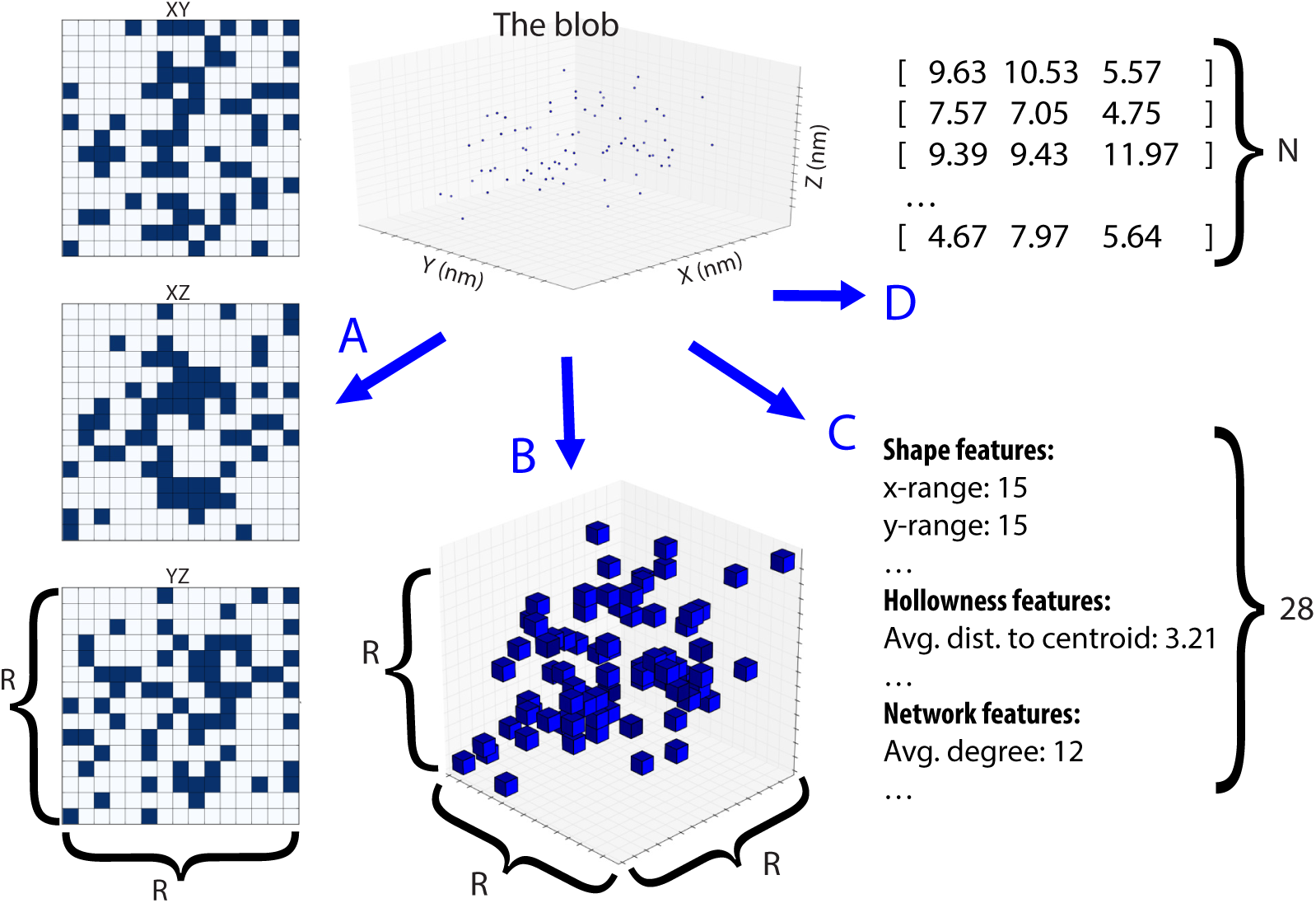
Possible input representations of a point cloud with range *R* and number of points *N*. (A) Three projections of the 3D point cloud onto 2D planes. Requires 3×*R*^2^ values to represent. This data representation is used for MVCNN. (B) A voxel representation. Requires *R*^3^ values to represent. (C) Hand-crafted/designed features. Requires 28 values to represent. This data representation is used for random forest. (D) A point cloud. Requires 3×*N* values to represent. Where *N* is the number of points per blob. This data representation is used for PointNet.

### Input (x)

Our SMLM dataset is 3D, i.e. contains location information for each molecule in all three dimensions. While the extra dimension provides additional information, which can improve the analysis of the data, three dimensional data also poses a number of possible pitfalls if one is not careful with how it is represented. The first is the size of the data.

The first versions of SMLM were only two dimensional, and therefore images can be neatly represented on a plane divided into pixels. If we expand this idea into three dimensions by dividing a 3D area into voxels, we get an exponential increase in size. Since the maximum range of our data is 512 nm, using 1 nm as our subdivision unit, an increase from 2D to 3D increases the size of a single blob from 262 thousand 2^18^ pixels to 134 million 2^27^ voxels. The second pitfall is the sparsity of each input data. The largest number of points belonging to a single blob is 512 points. If we encode this data in a 2D plane such that each point is encoded as a pixel with a value of 1 and every other pixel is has a value of 0, the ratio of effective bits (non-zero) is 2^9^*/*2^18^ = 0.2%. Expanding this to three dimensions and the ratio drops to 2^9^*/*2^27^ = 3*e* − 4%. From the above, it is clear that a voxel representation is ill-suited for the task at hand. Instead, we represent the data in three ways that avoid the above pitfalls.

i. **Expert features:** Relies on a simple analysis of the blob to generate hand-designed features reducing the input down to a size of 28 floating point numbers (Fig 4C).
ii. **Multi-view:** Transforms the 3D point cloud by projecting it onto three orthogonal 2D planes forming three 512*×*512 arrays of pixels (Fig 4A,B).
iii. **Point cloud:** Keeps the original point cloud representation from SMLM. When stored as a set of points, the data ratio of effective bits is 100%, and has a size of a number of points (512) *×* number of dimensions (3) (Fig 4D).

### Output (y)

We defined the output to be a one-hot encoding of the two classes, i.e. *y* = [1, 0] for positives, and *y* = [0, 1] for negatives. The two deep learning models (MVCNN and point cloud – PointNet below) first find a set of representative features *x*′ → *h*(*x*′) = *X*′, which are then linearly combined and passed through a softmax function *X*′ → *σ*(*w*^*T*^ *X*′+ *b*) = *ŷ*, where *w* is a learned set of weights and *b* is learned bias. From this it follows that *x* → *f* (*g*(*x*)) = *σ*(*w*^*T*^ *h*(*g*(*x*) + *b*). This approach significantly outperformed using a sigmoid to output a single number between 0 and 1.

### Different machine learning ML models

We have developed three models to best match the input representation. The deep features are the non-hand-crafted learned features extracted using the deep layers from either CNN or PointNet architectures. The hand-crafted features are the manually-designed features extracted based on previous domain knowledge [34].

#### Expert features – random forest classifier

The first model relies on 28 hand-crafted features that were chosen to capture different properties of the blobs based on known biology (Fig 4C). The 28 features describe the size (volume, XYZ range), shape (spherical, planar, linear), topology (hollowness), and network measures (degree, modularity, characteristic path, etc.) of each individual blob. To extract the shape features, we represented each blob as 3D point cloud centered at the blob mean of the points positions. Then, we used the eigendecomposition of the *N* × 3 matrix of every blob (Fig 4D) to extract the eigenvalues associated to the eigenvectors of the 3D matrix representation using the principal components analysis PCA method. The extracted eigenvalues are used to extract the different shape features of the blob. We mainly extracted the planer, linear, spherical, and fractional anisotropy (FA) shape features of every blob [35]. The volume is calculated using the convex hull of the Delaunay triangulation of the 3D matrix of the blob (Fig 4D). The hollowness features are extracted from the distance to centroid of the blob. We calculated the minimum, maximum, average, median, and the standard deviation of the distances from every point to the centroid of the blob. To extract the network features for every blob we represented the blob as a network where the nodes represent the points and the edges represent the proximity between every pair of nodes. We picked the proximity threshold for the network construction such as every blob in our dataset is one connected component. Then, the network features [36] are extracted from the constructed network for every blob [9]. The final feature vector is composed of all the extracted features and has a dimension of 1 × 28 [9] (Fig 4C).

We adopted machine learning random forest classifier [37] trained on the 28 hand-crafted features to automatically identify the blobs. Additionally, our goal is to design a machine learning classification model that generalizes well and therefore could be used to classify blobs not seen by the model. However, overfitting and underfitting cause poor performance and might prevent the model from generalization. To generalize better and avoid overfitting in our model, we used the bagging. Specifically, we leveraged Matlab TreeBagger toolbox. TreeBagger trains a large number of strong learners (i.e. random forest trees) in parallel. Then, it combines the results of all the trees to smooth out their predictions.

To evaluate the performance of the classifier to identify positive and negative classes of the blobs, we used the binary classification evaluation measures. Specifically, we used accuracy, sensitivity, and specificity measures. After the classification process, we need to count: the number of correctly identified blobs from the positive class which is known as true positive (TP), the number of correctly classified blobs from the negative class which is known as true negative (TN), the number of incorrectly identified (i.e. misclassified) positive blobs which is known as false negative (FN), and number of misclassified negative blobs which is known as false positive (FP). The classification accuracy is a statistical measure used to assess the performance of the binary classifier in identifying the number of correctly classified blobs to the total number of the examined blobs. Formally, 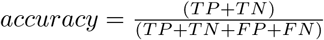. The sensitivity measures the ability of the classifier to correctly identify the positive blobs. Formally, 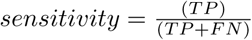. The specificity, on the other hand, measures the ability of the classifier to correctly identify the negative blobs. Formally, 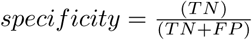.

Firstly, we need to validate the first model on the simulated dataset. We extracted 28 hand-crafted features for the simulated blobs from both classes (i.e. class 1/positive and class 2/negative). The random forest classifier is used to classify the blobs with number of trees equal 100. We used the TreeBagger Matlab implementation for the random forest. To evaluate the performance of the classifier on the simulated dataset, a 10-fold-cross validation is used. We obtained a 98.8% classification accuracy. The obtained specificity is 99% and sensitivity is 98%, which shows that the classifier can recognize the blobs from both classes with very low misclassification even when the number of points per blob is overlapping in both classes (Fig 3B). This shows the robustness of the used hand-designed features in identifying the blobs.

We then trained a random forest (RF) classifier using 100 trees in Matlab based on the extracted features from all the blobs in the dataset and using the binary labels of every blob. A 10-fold cross-validation is used to evaluate the classification results as seen in the first row of Table 1. A leave-one-cell-out is used in another experiment to evaluate the classification results also as shown in the first row of Table 2.

**Table 1.** Test set results on mixed cells.

|  | Accuracy | Sensitivity | Specificity | Parameters | Training time* |
| --- | --- | --- | --- | --- | --- |
| Hand-crafted | 0.92 | 0.97 | 0.86 | N/A | N/A |
| Multi-view | 0.92 | 0.98 | 0.85 | 31093187 | ~3500s |
| Point cloud | 0.81 | 0.79 | 0.83 | 3470284 | ~680s |

Results using sets made with blobs from each cell. \*Trained until convergence on a GeForce GTX 1060 GPU.

**Table 2.** Test set results using segregated cell.

|  | Accuracy | Sensitivity | Specificity |
| --- | --- | --- | --- |
| Hand-crafted | 0.94 | 0.97 | 0.90 |
| Multi-view | 0.94 | 0.98 | 0.90 |
| Point cloud | 0.83 | 0.72 | 0.96 |

Results using sets where each set uses different cells.

#### Multi-view – CNN

The second model applies a Convolutional Neural Net (CNN) to 2D multi-view data representations with projections of the point clouds onto 3 planes (xy, yz, xz) as shown in Fig 4A. Following Su et al. [38] naming convention, we call this model Multi-View CNN (MVCNN). A straightforward CNN architecture using alternating layers of convolutions and pooling and two final fully connected (FC) layers worked well. Variations to this model showed no discernible improvement. The layers of the CNN are as follows (Fig 5): conv1 (3×32), pool1(3×3), conv2 (32×64), pool2 (3×3), conv3 (64×128), pool3 (3×3), conv4 (126×256), pool4 (3×3), conv5 (256×512), pool5 (3×3), FC (256), FC (512), FC (2). A ReLu activation function was used on every layer except for the final fully connected layer, which uses a softmax activation. A cross entropy loss was used for the objective function, with the addition of a L2 weight regularization term.

**Fig 5.**
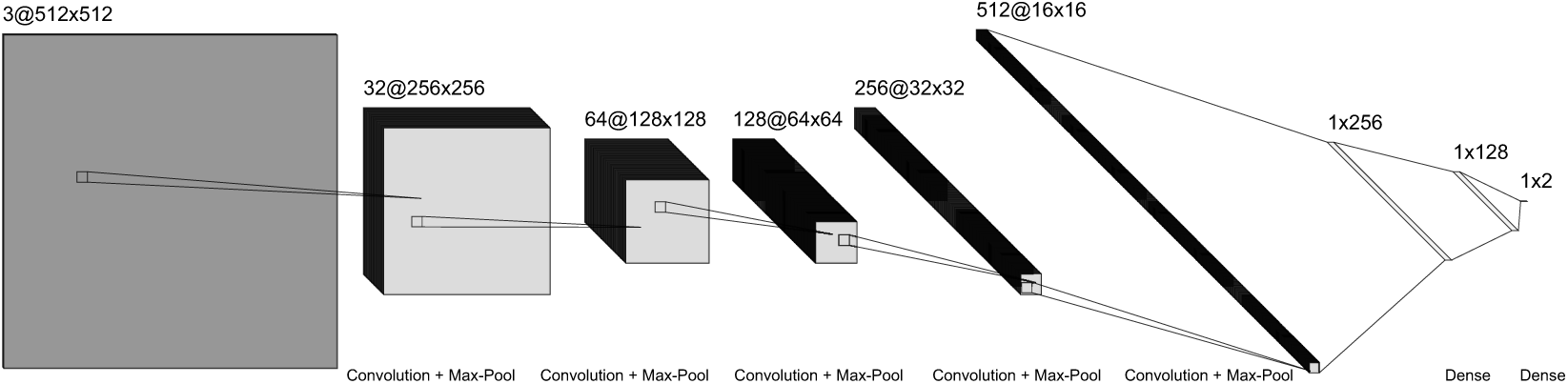
The architecture of the network used in the MVCNN. The layers of the CNN are: conv1 (3×32), pool1(3×3), conv2 (32×64), pool2 (3×3), conv3 (64×128), pool3 (3×3), conv4 (126×256), pool4 (3×3), conv5 (256×512), pool5 (3×3), FC (256), FC (512), FC (2)

#### Point cloud – PointNet

The third model is based on PointNet, which takes as input a set of 3D points. Minimal changes were made to the model described in [39]. In summary, PointNet uses the symmetric max function to enable its input to be unordered, as in the case of a point cloud. A number of hidden layers are used before the max function to transform the points into a higher dimensional space. The output of the max function is a representation of the point cloud and is passed through an FC network to classify the blob. For more detail see [39]. The alterations made are the removal of the dropout layer and of the data jittering, both of which were found to lower results. Consistent with the MVCNN model, a cross entropy loss was used. Qi et al. [39] demonstrates the success that both the MVCNN and PointNet approach can have on point cloud classification.

### Evaluation Methodology

To evaluate our model, we divide the 1714 blobs (the positive and the sampled negative blobs) into a training set, a validation set, and a test set in two different ways. The first way of creating the sets involves mixing the blobs of each cell, then keeping 200 blobs as a test set, using 100 blobs as a validation set, and using the remaining 1414 blobs as a training set. The second way is keeping cell 1, containing 124 blobs, as a test set, using cell 2, containing 100 blobs, as validation, and using the remaining 1290 blobs from the other cells as a training set. Each of the above sets is balanced in terms of negative and positive blobs. The use of the two groupings reveals if the data from one cell can be generalized to other cells.

### Mixed blobs

From the above results, we see that the hand-designed features and multi-view models generate similar results, while the point cloud model falls behind. A fundamental difference between the point cloud input and the other inputs is that it is un-ordered i.e. a blob can be mapped to more than one representation. The hand-designed features have a human chosen order. The multi-view input is a projection of the data on a 2D plane, which forces the data into a geometrical ordering. In point clouds, however, changing the order of the points does not change the underlying blob. The results would support the hypothesis that a useful order to data benefits data analysis.

While it does perform worse on the primary metrics, it is important to note that the point cloud input does have some advantages. First, compared to the hand-designed features, it does not require any preliminary analysis or expert knowledge. Second, compared to multi-view, the input data size and number of parameters is significantly smaller, and consequently, the model trains significantly faster. Finally, if segmentation of caveolae was a concern, both hand-designed features and multi-view would encounter major obstacles, but it has been demonstrated in [39] that it is possible to segment point clouds using PointNet.

### Cell-wise blobs

From the cell-wise results, we can show that knowledge learned can be generalized to other cells. This is important as it demonstrates the usefulness of this model on unlabeled blobs from future cells. The small increase in performance could be due to the slightly larger training set, or simply that the randomly chosen test cell contained an easier set of blobs to identify.

In both tables, the multi-view and hand-designed features approaches performed similarly well. However, we believe that an increase in dataset size may be more beneficial to the deep learning approach, meaning that using a larger dataset may allow the multi-view approach to outperform the hand-designed features. As we continue to collect more data, we hope to test on a larger dataset in the future to confirm this hypothesis.

The higher sensitivity (in both Table 1 and Table 2) suggests that our learned models are capable to identify the caveolae blobs more accurately, whereas the relative lower specificity means that our learned models are less accurate in identifying the scaffolds. This opens the door for further study of the scaffolds and suggests that those biological structures are more complex and have higher variation than the positive blobs. We expect more than one sub-category in the negative blobs. Moreover, the negative blobs in PC3 population might be different from the negative blobs in PC3-PTRF population (i.e. the CAVIN1/PTRF might also affect the structure of the scaffolds). We leave this investigation for the future as it requires more biological experiments and data.

### Hand-crafted/designed VS. deep features

Multiple data representations have a critical impact on the performance of the final semantic learning task. For classification task, the separability of the classes is highly dependent on the features and the way they were extracted. Fig 6 shows the t-SNE visualization of the features where the high-dimensional feature space is projected onto a 2-dimensional space [40]. The hand-crafted and MVCNN features are more clustered and separable compared to the PointNet features. However, the classes in this 2D projected view are not perfectly separable. This is likely due to the negative class having many complex subcategories, which depicts the complexity of the classification tasks at hand.

**Fig 6.**
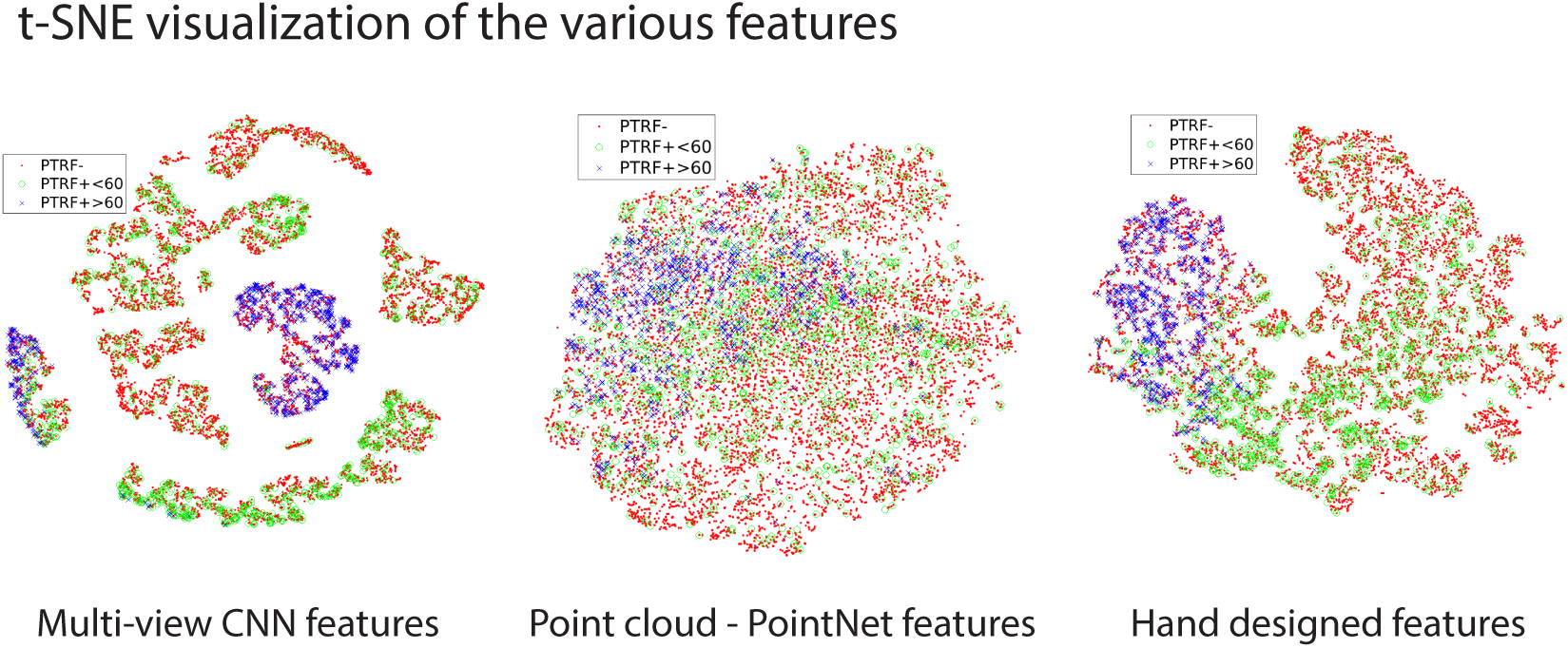
2D t-SNE visualization of the projected feature space from the various data representations used for identifying the Cav1 blobs. We visualize the projection of high-dimensional feature space into 2-dimensional space using t-SNE from the MVCNN, point cloud – PointNet, and hand-crafted features. Every point in the t-SNE plot represents the projected features of a blob. The red, green, and blue points represent the projected features of the PTRF-, PTRF+< 60, and PTRF+*≥* 60 blobs respectively.

The trade-offs (Table 3) between the different methods used to represent and classify the blobs in this work involve time and space (memory) complexity of training and inference, classification accuracy achieved, interpretabilty of the discriminant features, and the level of automation required (amount of human involvement). See Table 1 for the time and computational complexities of the deep learning methods.

**Table 3.**
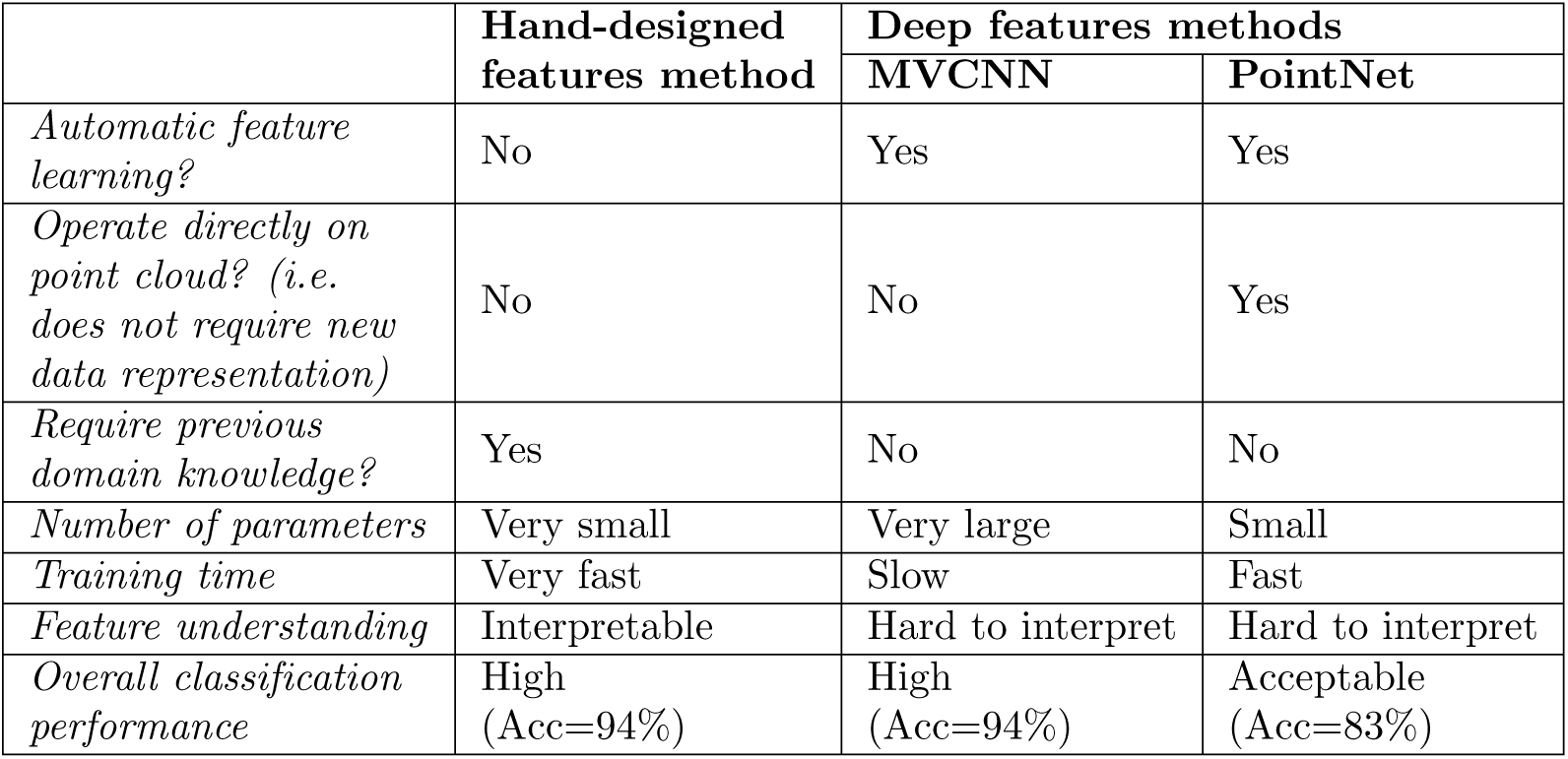
Hand-crafted and deep (MVCNN and PointNet) features trade-offs.

The key advantage of deep learning is that it avoids the manual process of constructing and selecting hand-designed and engineered features and that it boasts fast inference. However, the requirement of large training dataset, large computational resources for training, and its opaque uninterpretable, black box models are still major issues in deep learning.

Deep learning approaches that operate directly on unstructured data, such as PointNet that consumes the point cloud directly without any transformation, have the additional advantage of retaining the compactness and precision of the original data.

We hypothesize that the inferior classification accuracy performance of PointNet is due to its unordered input. PointNet was originally tested using a dataset that is an order of magnitude larger than ours, and it is possible that with a larger dataset the model would be able to learn to overcome the unordered nature of its input.

MVCNN capitalizes on the highly successful CNNs to achieve superior performance in classification accuracy but at the expense of longer training times and requiring large underlying representations, i.e. a large number of small pixels, needed to diminish quantization errors (compared with the pure 3D point cloud input adopted by PointNet).

Albeit being easily interpretable (which MVCNN and PointNet are not) and achieving higher accuracy than PointNet, hand-crafted features used in conjunction with classical machine learning approaches (e.g. RF) require prior expert knowledge of the biological structures in order to design and select features, which is may not always be feasible especially in scientific discovery. We summarize the trade-offs of the hand-crafted and deep features in Table 3.

## Conclusion

Our research into the analysis of super-resolution images using machine learning algorithms has yielded a number of successful techniques that can be used to accurately and automatically predict whether or not a blob is a caveola. Both using hand-designed features, as well as applying a convolutional neural net to projections of the point cloud, performed similarly well while using PointNet on a point cloud was less successful. Classifying biological structures at the cell membrane is of importance as it allows the biologist to study the relationship between structure and function. It could also be used to identify biomarkers for the different structures that could enable drug design at the molecular level and potentially lead to disease therapy.

## Future work

Further research on this topic would greatly benefit from additional labelled data. SMLM data for both PC3 and CAVIN1/PTRF from the same labeled cell would provide additional and more precise labels than the current method which relies on a wide-field TIRF CAVIN1/PTRF mask of lower resolution. Additional data would include double labeled SMLM images with high-resolution localizations for both Cav1 and CAVIN1/PTRF that would provide us with a more accurate class blob label. Moreover, the proposed methods described in this paper could be applied to other applications and other labeled proteins to automatically characterize the underlying biological structures. The feature extraction either via hand-designed or automatically derived features via deep learning could be applied to any SMLM data after extracting the SMLM clusters for the different machine learning tasks. We applied our method to Cav1 protein clusters from SMLM images. However, the methods are applicable to other SMLM biological data/applications.

While the current methodology relies on binary classification, caveolae or not-caveolae, it is likely that the not-caveolae class may be better represented as many classes. Using unsupervised methods such as k-means or mixture of Gaussians can allow us to subclassify the non-caveolae structures into more representative classes [9]. Applying similar models to ones described in this paper to a multi-class version of the problem may increase performance if the classes are better a representation of the true data.

Future work could also involve examining methods for interpreting deep learning models (e.g. [41]) applied to biological structures, and exploring research trends in unsupervised deep learning. It will also be interesting to explore developing deep neural network layers from the ground up particularly targeted to processing typical visual patterns seen in biological structures (as opposed typical man-made objects common in computer graphics applications).

## Acknowledgments

We thank Dr. Keng C Chou (Chemistry, UBC) for helpful comments and discussion.

## Notes

#### Summary of Updates

Figures are embedded in the manuscript in this revision. A new simulated data is added with a discussion (new Fig. 3 is introduced as well). The mathematical definition of the evaluation measures is added. Update Table 1 with time and computational complexity. Add new and related references. Clarify some terms in the paper.

https://doi.org/10.6084/m9.figshare.7932326

